# Novel high-resolution ion mobility mass spectrometry for site-specific quantification of the sirtuin-5 regulated kidney succinylome

**DOI:** 10.1101/2025.11.01.686011

**Authors:** Birgit Schilling, Leonard Rorrer, Lauren Royer, Anne Silva-Barbosa, Katherine Pfister, Eric Goetzman, Sunder Sims-Lucas, Daniel DeBord, Joanna Bons

**Affiliations:** Buck Institute for Research on Aging, Novato, CA, USA; MOBILion Systems, Inc., Chadds Ford, PA 19317, USA; Department of Pediatrics, University of Pittsburgh Medical Center Children’s Hospital of Pittsburgh (UPMC CHP), University of Pittsburgh, Pittsburgh, PA 15224, USA

## Abstract

Protein post-translational modifications (PTMs) dynamically regulate essential biological and cellular processes. Lysine succinylation changes the amino acid charge, potentially affecting protein structures and functions, and dysregulation of protein succinylation may lead to metabolic disorders. Proteome-wide succinylation quantification using proteomic tools remains challenging, especially due to the low abundance of succinylated peptides and the frequent presence of isomeric PTM forms. Ion mobility spectrometry workflows that can differentiate peptidoforms with different PTM distributions represent a powerful strategy to alleviate these challenges. Recently, a new Parallel Accumulation with Mobility Aligned Fragmentation (PAMAF™) operating mode for high-resolution ion mobility-mass spectrometry (HRIM-MS) analysis based on the structures for lossless ion manipulation (SLIM) technology was introduced. Here, we first assessed the performance of PAMAF mode for protein succinylation analysis using synthetic succinylated peptides, demonstrating residue-level differentiation of co-eluting isomers and isobars and precise PTM site localization. We leveraged this novel approach to investigate succinylome remodeling in kidney tissues from wild-type and Sirtuin-5 (Sirt5) knock-out mice, a NAD^+^-dependent lysine de-succinylase. PAMAF acquisitions yielded ∼1,000 confidently identified and accurately quantified succinylated peptides and sites from mouse kidney. Sirt5 regulated succinylation of mitochondrial proteins involved in metabolic processes, including fatty acid oxidation, the tricarboxylic acid cycle, and propionate metabolism.

## Main text Introduction

Protein post-translational modifications (PTMs) are critical regulators of protein conformations, activities, interactions, and subcellular localization, and they dynamically modulate a plethora of cellular signaling processes and biological functions. Lysine residues can be subjected to multiple types of modifications, including the recently discovered succinylation^1^. This reversible PTM profoundly alters lysine properties by increasing the size of its side chain and reversing its charge from +1 to -1 under physiological conditions, eventually leading to significant conformational and functional changes to the protein^2^. Succinylation occurs when a lysine residue reacts with the donor metabolite succinyl-CoA, and the succinyl moiety becomes covalently attached to the primary amine. This modification is reversible and the removal of the succinyl group is dynamically regulated by lysine deacylases, including sirtuins. More particularly, sirtuin-5 (Sirt5) exhibits NAD^+^-dependent lysine de-succinylase activity^3^. Alterations of protein succinylation profiles have been linked to various pathological conditions including metabolic disorders, neurodegenerative diseases, cardiovascular diseases, and cancers^4–6^.

Mass spectrometry (MS) has become the method of choice for large-scale, proteome-wide PTM profiling and quantification. Significant advances in succinylated peptide enrichment strategies, liquid chromatography coupled with tandem mass spectrometry (LC-MS/MS) methodologies, and computational pipelines have uncovered thousands of distinct succinylation sites in various biological systems^7–13^. Despite these major improvements and the high relevance of succinylation biology, a major analytical barrier remains to fully appreciate the dynamic and site-specific regulation of protein succinylation: the ability to differentiate positional succinylated isomers, *i.e.* a succinyl modification that occurs on the distinct lysine residues within the same peptide sequence or peptidoform. While the specific PTM sites may have distinct functional outcomes, conventional HPLC-MS/MS proteomic strategies are typically limited in resolving isomeric species that may co-elute, and share the same mass-to-charge (m/z) ratio, thus generating chimeric/overlapping fragmentation MS/MS spectra. Subsequently, HPLC and MS dimensions are often insufficient to fully profile and accurately quantify lysine succinylation. Subsequently, our understanding of the dynamic and site-specific regulation of succinylated proteoforms remains elusive.

Interfacing ion mobility (IM) spectrometry technology with high-resolution mass spectrometers offers an orthogonal approach to chromatographic and *m/z* separations. Ion mobility separates peptide ions based on their size, shape, and charge through their migration in an inert gas under an applied electric field. Various IM techniques, including trapped ion mobility spectrometry (TIMS)^14^ and high-field asymmetric waveform ion mobility spectrometry (FAIMS)^15^, have been successfully employed for large-scale PTM analysis, demonstrating improved structural isomer differentiation compared to conventional HPLC-MS/MS workflows. However, ion mobility resolution and duty cycle achieved still present limitations. The structures for lossless ion manipulations (SLIM) technology, developed at PNNL, implements traveling wave ion mobility (TWIM) and uniquely enables ion packets to be shuttled around “corners”^16, 17^. This allows for long serpentine ion path geometries that increase the mobility path length within a reasonable form factor device, enhancing resolving powers across the full mass spectrum. Potentially this enhanced ion mobility separation may greatly improve the reduction of complexity of proteomes and lead to improved quantification of all analytes. In this study, we employed a high-resolution ion mobility (HRIM) system developed by MOBILion Systems that incorporates a 13 meter serpentine SLIM ion path interfaced with a high-resolution quadrupole-time-of-flight (Q-TOF) mass spectrometer, achieving peak resolving powers above 200 in a single pass^18^, for lysine succinylation analysis. While Chouinard *et al.* previously reported the coupling of HPLC and SLIM-MS for targeted analysis of selected synthetic phosphopeptides at the MS1 level only^19^, more recent work by Arndt *et al.* applied HPLC-HRIM-MS with collision-induced dissociation (CID) for targeted PTM analysis on monoclonal antibodies for biopharmaceutical assays^20^. In our study presented here we performed unbiased discovery / non-targeted analysis of PTM peptides by SLIM-MS. Here, we leveraged the novel Parallel Accumulation with Mobility Aligned Fragmentation (PAMAF™) operating mode using a modified MOBIE^®^ system that enables precursor and fragment ion analysis without any ion filtering by the quadrupole, for proteome-wide succinylome analysis. More particularly, we demonstrated for the first time the application of LC-PAMAF-MS to confidently differentiate succinylated isomers using synthetic succinylated peptides and to accurately quantify hundreds of distinct succinylation sites from complex Sirt5- deficient (Sirt5^-/-^) and wildtype (WT) mouse kidney extracts.

## Results

### Parallel Accumulation with Mobility-Aligned Fragmentation (PAMAF)-mass spectrometry for protein succinylation analysis

A modified MOBIE system operated in PAMAF mode was used for protein succinylation analysis via two experimental paradigms (**Fig. 1A**). In study S1, a mixture of heavy-labeled synthetic succinylated peptides and non-modified counterparts were analyzed to assess PAMAF operation performance for confident PTM identification and site localization as well as succinylated isomer differentiation. In study S2, we expanded our analysis to examine a more complex matrix, namely enrichments of proteolytic succinylated peptides derived from mouse kidney tissues, to investigate the dynamic remodeling of the succinylome upon Sirt5 deletion, a NAD^+^-dependent lysine deacylase with de-succinylase activity^3^.

**Figure 1.**
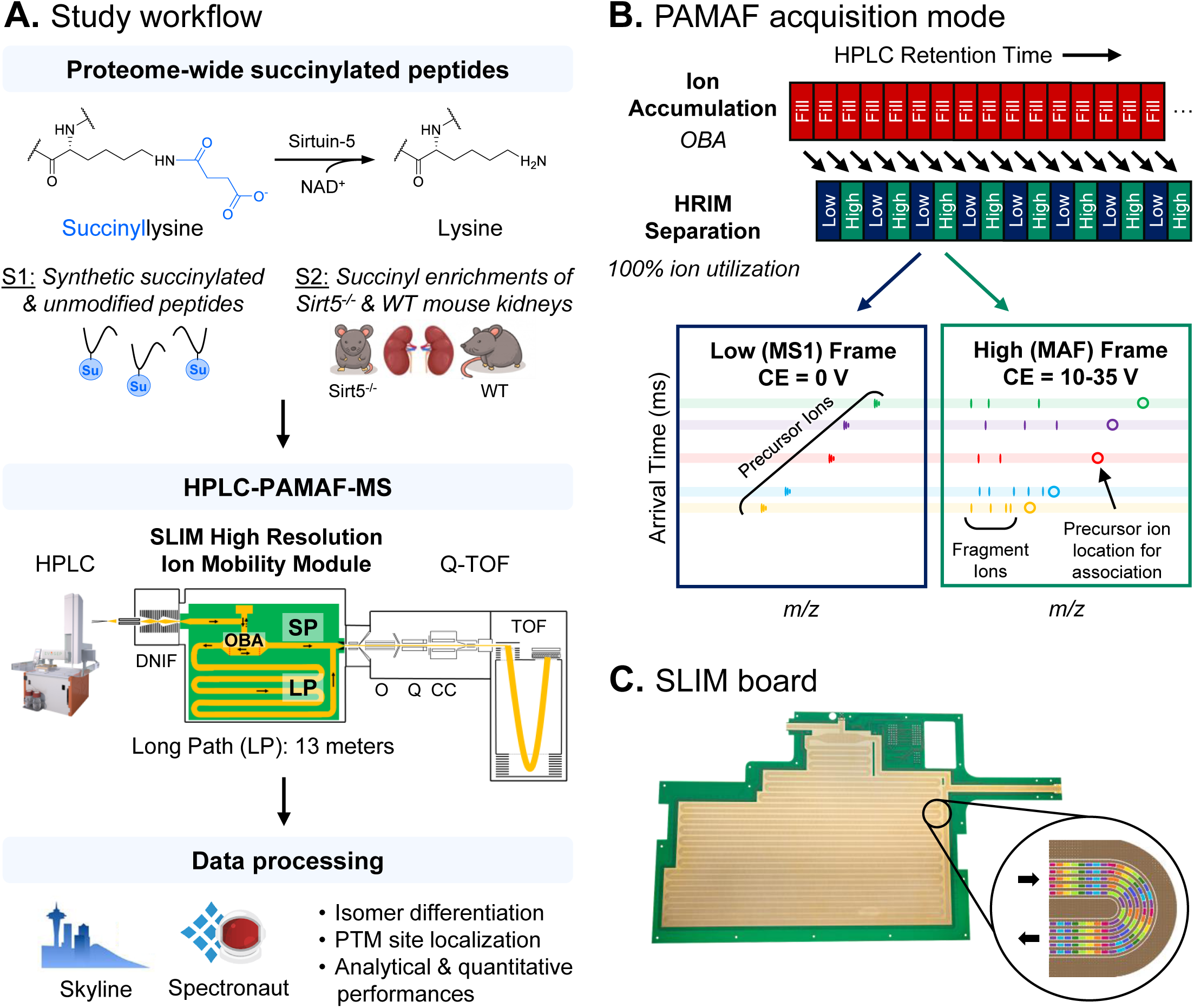
Experimental design for proteome-wide succinylomic analysis using the Parallel Accumulation with Mobility-Aligned Fragmentation (PAMAF) mass spectrometry technology. A. To assess PAMAF mode performance for protein succinylation identification and site localization, a first study (S1) on synthetic mouse succinylated and matched unmodified peptides was conducted. Then, PAMAF technology was applied to a second study (S2) to investigate the dynamic mouse kidney succinylome remodeling upon Sirtuin-5 (Sirt5) deletion, a NAD^+^-dependent lysine deacylase with de-succinylase activity. Data were collected on a prototype HPLC-QTOF platform adapted from the MOBILion Systems MOBIE device, equipped with a Structures for Lossless Ion Manipulation (SLIM)-based high-resolution ion mobility (HRIM) device, and analyzed with Skyline or Spectronaut. B. The PAMAF mode consists of alternating low and high collision energy (CE) mobility frames containing precursor (MS) and fragment (MS/MS) ions, respectively, throughout the chromatographic gradient. Ion capacity is maximized through parallel ion accumulation in the onboard accumulation (OBA) region prior to HRIM separation. Finally, precursor and fragment ions are correlated based on arrival time (AT) alignment in the IM dimension. C. Cross-sectional view of the SLIM board that uses a 13-m-long serpentine path (long path) for traveling wave ion mobility spectrometry. AT, Arrival time. CC, Collision cell. DNIF, Dual narrow ion funnel. HRIM, High-resolution ion mobility. LP, Long path. MAF, Mobility-aligned fragmentation. O, Octopole. OBA, onboard accumulation. PAMAF, Parallel Accumulation with Mobility-Aligned Fragmentation. Q, Quadrupole. SLIM, Structures for Lossless Ion Manipulation. SP, Short path. TOF, Time-of-flight.

After liquid chromatographic separation, precursor ions entered the entered the HRIM module where ion mobility is performed using the SLIM assembly, which is connected online to an Agilent 6545 Q-TOF mass spectrometer for detection. More specifically, the ions entered the HRIM module, where they were first accumulated in the onboard accumulation (OBA) region for 100 ms, followed by high-resolution ion mobility (HRIM) separation in a long serpentine that is 13 meters long (long path, 400 ms)^18^ (**Fig. 1B-C**). The SLIM assembly within the HRIM module allows for parallel accumulation of ions prior to mobility separation, achieving ∼100% ion utilization (100% duty cycle). Ions leaving the SLIM module then pass through the quadrupole (Q) which was operated as an ion guide without any mass filtering, gating, or ion selection, thereby enabling capture of the entire m/z range for both MS1- and MS2-level analyses. The collision cell (CC) was operated alternately at low collision energy (CE; MS1 scan) or high CE for fragmentation (MS2 scan) as shown in **Fig. 1B**. Lastly, ions were detected by the high-resolution TOF analyzer, where a full mass spectrum was acquired for every mobility time point. After precursor ion accumulation and mobility separation, a low CE (MS1) frame recording *m/z* and arrival time (AT) features of all co-trapped precursor ions is followed by a high CE (MAF, MS2) frame capturing the ones of the fragment ions resulting from fragmentation of all precursor ions were recorded during each cycle. As precursor ions and their respective fragment ions share the same arrival time, they could be associated by AT alignment in the IM dimension.

Finally, collected data was processed with the open-source Skyline^21^ and Spectronaut (Biognosys) tools, that support PAMAF-MS data for succinylated peptide identification, succinyl site localization, and quantification.

#### Succinylation site localization of ion mobility-resolved co-eluting isomers

To assess the performance of PAMAF mode to precisely localize the PTM sites and confidently differentiate succinylated isomers, we used a well-controlled sample set of synthetic heavy-labeled peptides composed of 17 succinylated peptides, including four pairs of succinylated isomers, plus 13 matching non-modified peptides (total: 30 peptides; **Fig. 2A**, **Supplementary Table 1**). For each peptide in the mixture, 500 pg were injected on-column. Peptides were first separated by LC using the Evosep 200 samples per day (SPD) method (5.6-min gradient), then by HRIM in the MOBIE SLIM module for PAMAF mode analysis. Data were analyzed in Skyline, which supports the raw HPLC-PAMAF-MS data (*.mbi data) generated by the MOBIE system. Here, despite a very fast chromatographic gradient with peptide elution within ∼3.5 min (**Fig. 2B**), the extracted ion chromatograms (XICs) of the succinylated peptides were defined with nine points on average (ranging from six to 13 points across the chromatographic peaks; **Fig. 2C**). This is important as ultra-fast HPLC coupled to MS often yields a low number of data points across the chromatographic peaks, which can lead to inaccurate quantitative measurements.

**Figure 2.**
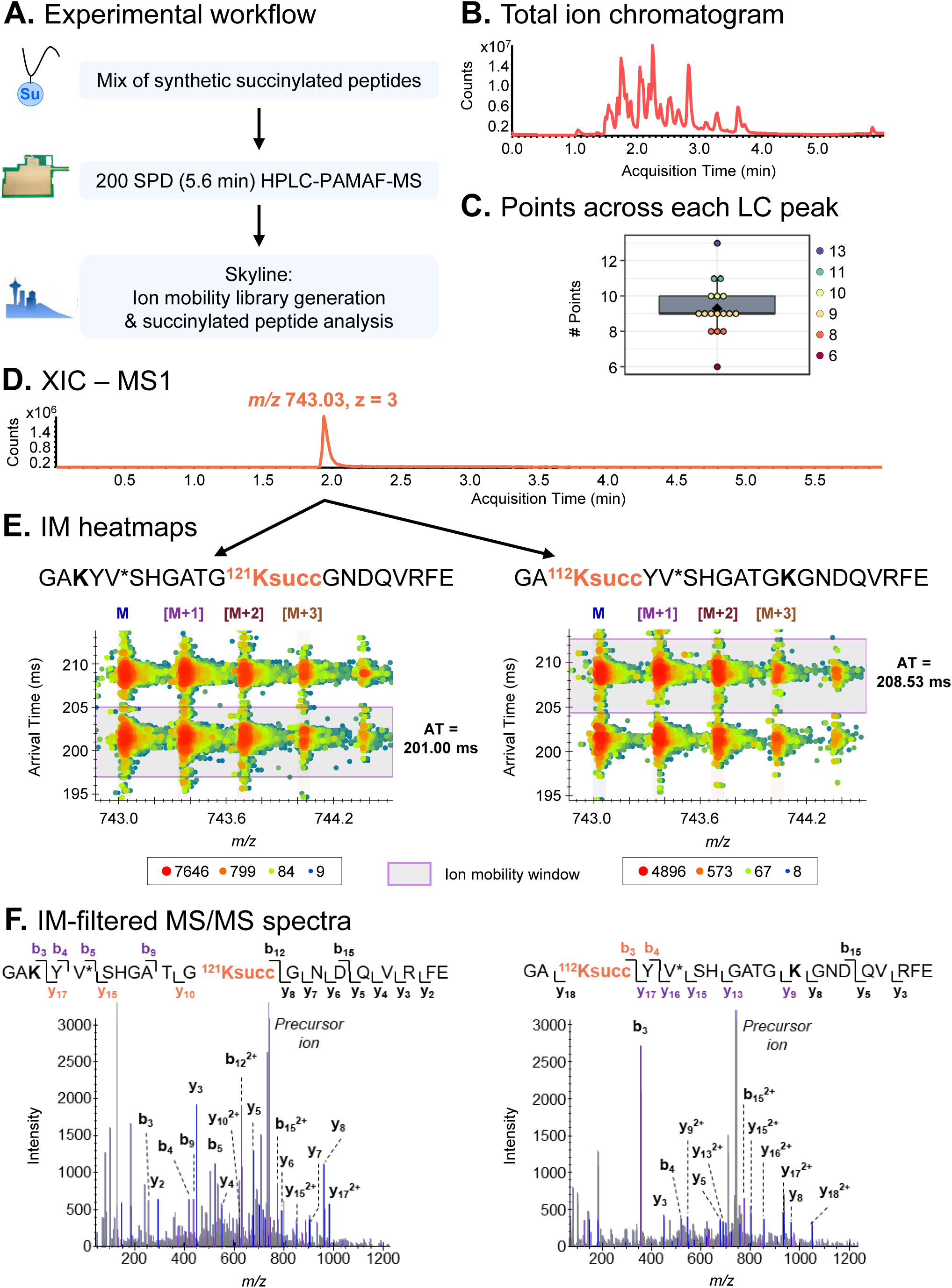
The PAMAF technology enables confident identification and site localization of co-eluting succinylated isomers. **A.** A mix of 17 synthetic mouse succinylated peptides, including 4 pairs of isomers, and the 13 matched unmodified peptides was analyzed on an Evosep UHPLC coupled to a prototype SLIM-based HRIM Q-TOF (Agilent 6545) MS operated in PAMAF mode. Skyline was then employed for ion mobility (IM)-aided feature extraction and succinylated peptide analysis. **B.** Representative total ion chromatogram. **C.** Distribution of the number of points across the peaks obtained for the succinylated peptides separated by a 5.6-min gradient. The black diamond represents the average value. **D-E.** The extracted ion chromatogram (XIC) of the precursor ion at *m/z* 743.03 (z = 3) shows one visible peak, while two species are detected and confidently separated by ion mobility. The purple windows on the heatmaps represent the ion mobility window defined in the Skyline ion mobility library for accurate IM-based filtering (integration band = 50 Rp). **F.** IM-based filtered MS/MS spectra for the two co-eluting species at *m/*z 743.03 (z = 3) corresponding to the succinylated isomers GAKYV*SHGATG**^121^Ksucc**GNDQVRFE (*left*) and GA**^112^Ksucc**YV*SHGATGKGNDQVRFE (*right*) from the mouse argininosuccinate synthase (UniProt ID: P16460). AT, Arrival time.

A typical challenge of IM-MS analysis is accurate data extraction and signal integration when applying mobility-based filtering – a requirement that becomes critical when analyzing isomeric or isobaric species. To overcome this, we first generated an assay-specific ion mobility library in Skyline to determine the arrival times for the peptides^22^ (**Supplementary Table 2**). To note, library arrival time values can be reviewed and manually adjusted if needed. The resolving power (Rp) was optimized to enable high specificity and selectivity of the mobility separation using narrow integration windows, while ensuring high accuracy by capturing the full mobilogram peak. Here, a fixed Skyline Rp setting of 50 appeared optimal for succinylated peptide analysis. As illustrated in **Fig. 2D-E**, a single chromatographic peak was observed when extracting the ion chromatograms for the two succinyl isomers GA**^112^Ksucc**YVSHGATGKGNDQVRFE and GAKYVSHGATG**^121^Ksucc**GNDQVRFE at m/z 743.03. These two co-eluting species could not be resolved by liquid chromatography, potentially leading to false or incomplete identifications. However, when applying mobility filtering, two distinct isotope distributions were detected in the heatmap: one at AT 201.00 ms and the other at 208.53 ms. Based on the AT values contained in the library and the defined Rp, mobilograms were integrated as represented by the purple IM window on **Fig. 2E**. Retrieved mobility-filtered MS/MS spectra showed very good sequence coverage with isomer-specific fragment ions, enabling confident peptide identification (**Fig. 2F**). More particularly, the detection of the ions y_10_, y_15_, and y_17_ containing the succinyl group (in orange for the sequence on the left) and ions b_3_, b_4_, b_5_, and b_9_ with no mass shift (in purple) confirmed the PTM site localization on the second lysine residue for the precursor ion arriving at 201.00 ms, *i.e.* Lys-121. In a similar but reversed manner, the succinyl group was localized on the first lysine (Lys-112) for the species at AT 208.53 ms. Another example of succinylated peptide isomers is presented in **Supplementary** Fig. 1. The mobility-filtered fragmentation spectra for LTGA**^216^Ksucc**VVVSGGRGLKSGE and LTGAKVVVSGGRGL**^226^Ksucc**SGE (precursor ions at m/z 607.67) contained numerous discriminating fragment ions (in blue and purple on the sequences) that supported the precise PTM localization site. The application of IM filtering for these two mobility-resolved species enabled confident isomer differentiation, even when shared fragment ions were extracted (e.g., ions b_4_ and y_16_).

Altogether, this demonstrates that HRIM using PAMAF mode is a powerful tool for differentiating co-eluting succinylated isomeric peptides that could not be resolved without ion mobility and to confidently localize their PTM site. In addition, this study highlights the compatibility of PAMAF mode with ultra-fast liquid chromatographic gradients, which are necessary to enable large-scale and high-throughput proteomic assays.

#### Combined PAMAF mode and library-free directDIA for analysis of the Sirt5- regulated succinylome

We expanded our evaluation of PAMAF mode to a quantitative succinylome analysis of a more complex, biologically relevant system to study Sirt5-regulated protein succinylation in mouse kidney tissues. Kidneys from male whole-body Sirt5-/- mice and littermate WT controls (four mice per group), aged 3 months, were harvested and homogenized, protein lysates were digested with trypsin, and finally succinylated peptides were enriched using pan-succinyl antibody-coated beads (**Fig. 3A**). Succinyl enrichments were analyzed by HPLC-MS using PAMAF mode. Data were processed with directDIA in Spectronaut, which supports ion mobility data from the modified MOBIE platform. We first assessed the performance of the PAMAF workflow for nontargeted, large-scale PTM analysis (**Fig. 3B-G**). Representative total ion chromatogram (TIC) from a Sirt5^-/-^ succinyl enrichment showed that peptides were eluted throughout the LC gradient (**Fig. 3B**). Efficient non-linear retention time (RT) and ion mobility regression was achieved in Spectronaut (**Fig. 3C**). Regarding ion mobility separation, precursor ions from the same Sirt5^-/-^ succinyl enrichment showed arrival times (AT) from 100 to 400 ms (**Fig. 3D**). Succinylated precursor ions specifically exhibited *m/z* values from 353 to 1468 and ion mobility AT from 134 to 388 ms (**Fig. 3F**). Very good IM calibration was achieved, as shown in **Fig. 3E**. Quantified peptides span a dynamic range of 5 orders of magnitude in this assay, among which succinylated peptide median quantity covered 3.9 orders of magnitude (**Fig. 3G**). Overall, 1094 succinylated peptides corresponding to 980 unique succinylation sites from 267 protein groups were confidently identified and quantified in Sirt5^-/-^ and WT mouse kidneys in this study (**Fig. 3H**).

**Figure 3.**
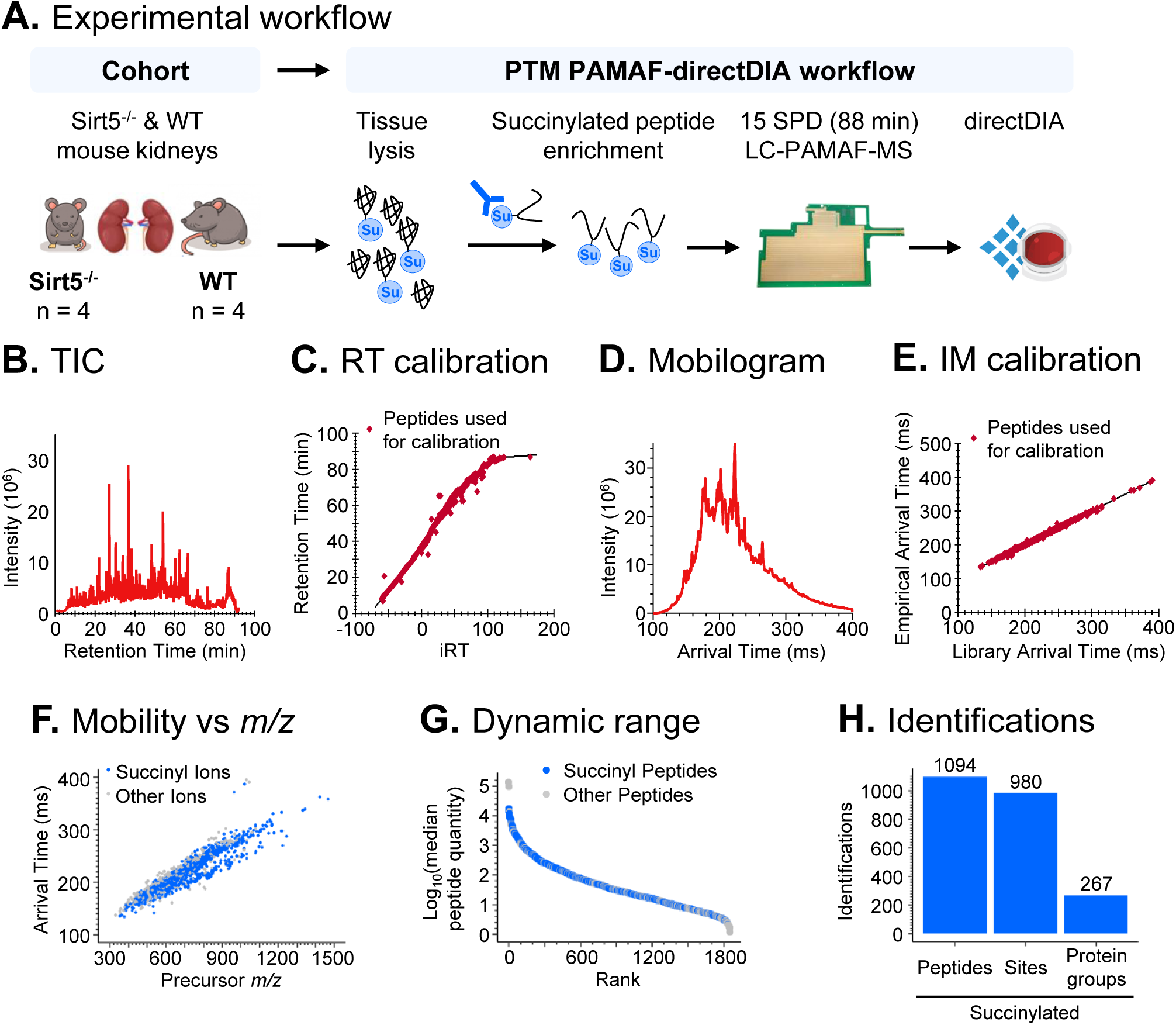
Evaluation of the PAMAF mode for complex mouse kidney tissue succinylomics. **A.** Kidney tissues were harvested from Sirt5^-/-^ and WT mice (n = 4 each) and lysed. After protein digestion, succinylated peptides were enriched using antibody-coated beads and analyzed on an Evosep UHPLC coupled to a prototype SLIM-based HRIM Q-TOF (Agilent 6545) MS operated in PAMAF mode. Collected data were analyzed using the spectral library-free directDIA software. **B,D.** Representative **B.** total ion chromatogram and **D.** mobilogram of a Sirt5^-/-^ succinylated peptide enrichment replicate. **C.** Retention time and **E.** ion mobility calibration of the Sirt5^-/-^ succinylated peptide enrichment replicate. The red dots correspond to the peptides used for the calibration and the black line to the regression curve. **F.** Distribution of the arrival time as a function of the precursor ion m/z. The blue dots correspond to succinylated precursor ions and the grey ones to non-succinylated ions. **G.** Dynamic range of the peptides quantified in the assay. Succinylated peptides are represented in blue. **H.** Number of identified and quantified unique succinylated peptides, sites, and protein groups. AT, Arrival time.

#### Differentiation of succinylated isomers from mouse kidneys

A central challenge in mass spectrometric analysis is the deconvolution of overlapping isotope patterns and the discrimination of isomeric and isobaric species – a complexity further exacerbated in large-scale, proteome-wide analyses of PTMs. For example, assessing the detectable precursor ions from *m/z* 799.9 to 803.7 within a PAMAF-MS1 spectrum (RT 64.44 – 64.82 min), two different isotope distributions were observed with the monoisotopic ions at *m/z* 800.66 and *m/z* 800.74 (**Fig. 4A**). The most intense precursor ion present in the mobilogram (AT 246.11 ms), MS1 spectrum (*m/z* 800.74), and heatmap zoomed displays was confidently identified as the succinylated sequence DYPLPDVAHVTMoxLSASQ**^60^Ksucc**ALK of the mitochondrial cytochrome c oxidase subunit 4 isoform 1 (COX41) with a succinyl group on Lys-60. Detected fragment ions containing the succinyl group provided direct evidence for the precise localization of the PTM site (in orange on **Supplementary** Fig. 2). Interestingly, the addition of SLIM-based HRIM separation helped to uncover the presence of a third species. Indeed, the heatmap illustrates that two distinct distributions were detected for *m/z* 800.66, that could not be resolved otherwise: one species at AT 257.43 ms and *m/z* 800.658 and the other one at AT 259.26 ms and *m/z* 600.661. The first released species (AT 257.43 ms) was further identified as the peptide TTGIVM^ox^DSGDGVTHTVPIYEGYALPHAILR of the cytoplasmic actin 1/2 (ACTB; ACTG), while the second released species (AT 259.26 ms) could not be identified. This demonstrates that HRIM using PAMAF mode achieved confident resolution of overlapping isotope distributions, uncovering isobaric species and peptidoforms.

**Figure 4.**
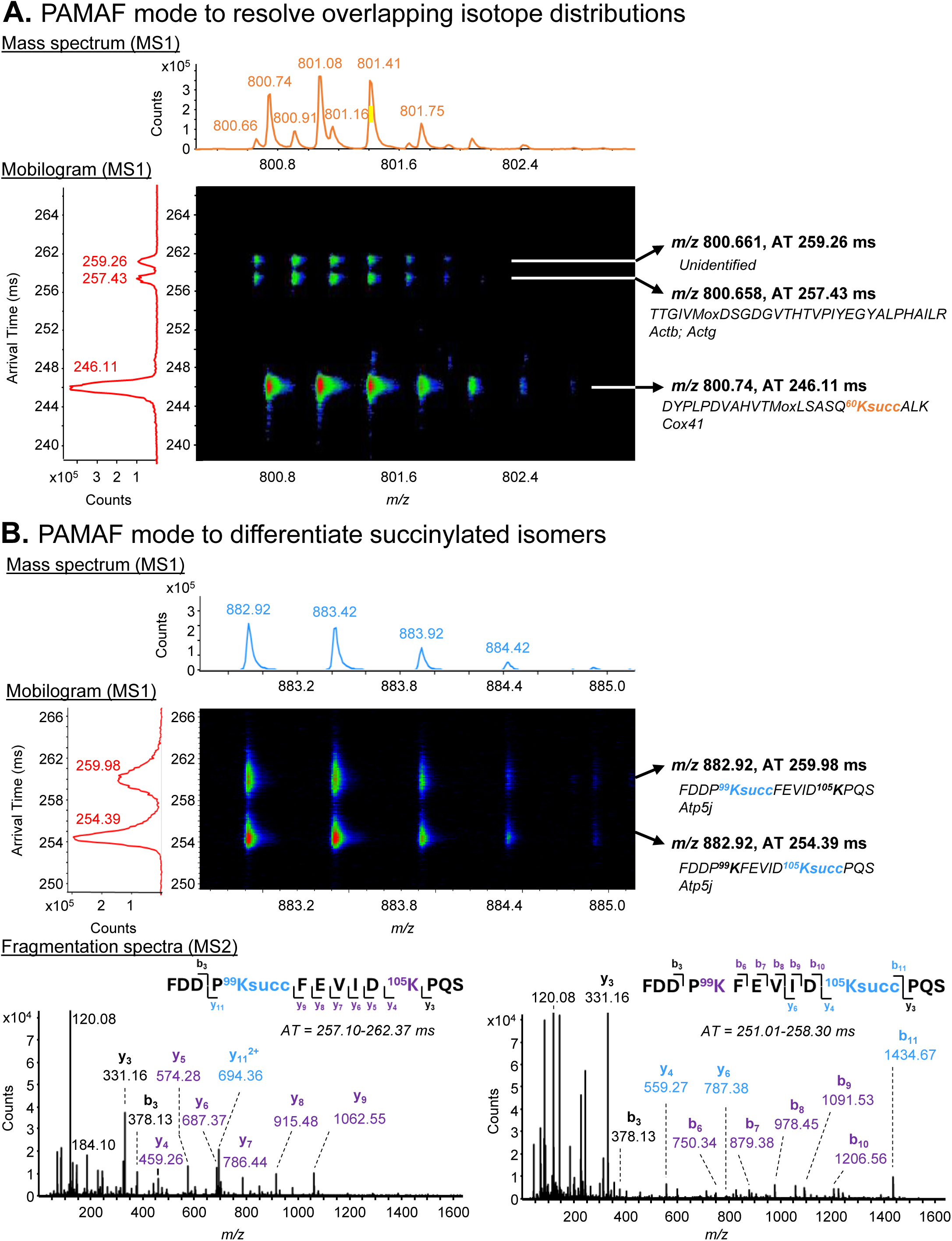
The PAMAF technology confidently resolves isotope distributions and differentiates succinylated isomers from mouse kidney tissues. **A.** Over the 64.44 – 64.82-min region, the zoom into the *m/z* range 799.9 – 803.7 of the MS1 spectrum (*top*) of a representative Sirt5^-/-^ succinyl enrichment shows two detected and distinct isotopic distributions with the monoisotopic ions at *m/z* 800.66 and *m/z* 800.74. The associated mobilogram (*bottom left*) reveals the presence of three distinct species as HRIM was able to differentiate two precursor ions with *m/z* 800.661 and arrival time (AT) 259.26 ms as well as *m/z* 800.658 and AT 257.43 ms. The most intense species as shown on the heatmap (*bottom right*) was *m/z* 800.74 and AT 246.11 ms that matched the succinylated peptide sequence DYPLPDVAHVTMoxLSASQ**^60^Ksucc**ALK of the mitochondrial cytochrome c oxidase subunit 4 isoform 1 (Cox4i1). **B.** Over the 56.69 – 57.80-min region, one isotopic distribution is detected on the mass spectrum (*top*) zoomed over the *m/z* range 882.8 – 885.0. The matching mobilogram and heatmap (*middle*) show the presence of two mobility-separated precursor ions at *m/z* 882.92 (z = 2). The MS/MS spectra (*bottom*) confirmed the identity of two succinylated isomers and their precise modification sites: FDDP**^99^Ksucc**FEVIDKPQS (*bottom left*; arrival time extraction window = 257.10 – 262.37 ms) and FDDPKFEVID**^105^Ksucc**PQS (*bottom right*; arrival time extraction window = 251.01 – 258.30 ms) from the mitochondrial ATP synthase-coupling factor 6 (Atp5pf). The MS/MS spectrum of the K-105 isomer is zoomed on the y-axis.

We then sought to assess PAMAF mode for succinylated isomer peptides with the same peptide sequence but alternate PTM localization sites. Precisely localizing modification site is critical, as different proteoforms may have different conformations, functions, and interacting partners. We were able to identify several pairs of succinylated isomers in mouse kidneys, among which were the peptides FDDP**^99^Ksucc**FEVID^105^KPQS and FDDP^99^KFEVID**^105^Ksucc**PQS with a succinylated lysine at position 99 and 105, respectively, of the mitochondrial ATP synthase-coupling factor 6 (ATP5J). At the *m/z* range corresponding to the doubly-charged precursor ion at *m/z* 882.92 on the PAMAF-MS1 spectrum (RT 56.69 – 57.80 min), a single isotopic distribution was detected (**Fig. 4B**). However, two species were distinctly separated by ion mobility, with base peak resolution, as observed in the MS1 mobilogram and heatmaps: one arriving at 254.39 ms and the other arriving at 259.98 ms. Reconstituted mobility-aligned fragmentation spectra for each species included discriminating fragment ions. More particularly, the presence of the ions y_4_, y_6_, and b_11_ with a mass shift corresponding to the addition of one succinyl group (+100.01 Da; ions in blue in the MS2 spectrum displayed on the right) but no succinyl group for the ion series b_6_ to b_10_ (in purple on the right spectrum) were direct evidence of the PTM localization site on the second lysine Lys-105 for the ion at AT 254.39 ms. Conversely, the presence of unmodified fragment ions y_4_ to y_9_ (in purple on the left MS2 spectrum) but ion y_11_ with a mass shift of 100.01 Da (in blue on the left spectrum) confirmed the presence of the succinyl group on the first lysine Lys-99 for the ion at AT 259.98 ms. Although separated by liquid chromatography (**Supplementary** Fig. 3), the additional and orthogonal IM resolution fully differentiated the two succinylated isomers, achieving highly confident identification and precise PTM site localization.

Altogether, this demonstrates that PAMAF mode achieved confident differentiation of isobaric species and succinylated isomer peptides in complex tissue samples, enabling in-depth site-specific succinylome profiling. This evaluation supported the investigation of the dynamic remodeling of the kidney succinylome in Sirt5^-/-^ and WT mice.

### Hypersuccinylation of mitochondrial metabolic proteins upon Sirt5 deletion

Supervised clustering using partial least-squares discriminant analysis (PLS-DA) on all identified and quantified succinylated peptides showed a genotype-based separation, that was at least partly explained by variate 1 (**Fig. 5A**). Comparison of the succinylation site relative abundance changes across Sirt5^-/-^ vs WT kidneys revealed that out of the quantified succinyl sites, 439 sites were significantly up-regulated and 22 sites were down-regulated (q-value <0.05 and |log_2_(Sirt5^-/-^ vs WT)| > 0.58; **Fig. 5B**, **Supplementary Table 3**). The top up-regulated sites were Lys-56 of the mitochondrial 10 kDa heat shock protein (CH10), Lys-265 of the mitochondrial acetyl-CoA acetyltransferase (THIL), Lys-147 of ADP/ATP translocase 1 (ADT1), Lys-90 of the mitochondrial ATPase inhibitor (ATIF1), and Lys-147 of the ADP/ATP translocase 2 (ADT2). Other mitochondrial Sirt5 targets included Lys-284 of the short/branched chain specific acyl-CoA dehydrogenase (ACDSB), Lys-209 of the 3-ketoacyl-CoA thiolase (THIM), Lys-392 of the methylmalonic aciduria type A homolog (MMAA), the ATP synthase-coupling factor 6 (ATP5J) with the four sites, Lys-46, Lys-84, Lys-99, and Lys- 105, as well as the cytochrome c oxidase subunit 4 isoform 1 (COX41) with Lys-60 and Lys-61. Some proteins were heavily targeted by Sirt5. For instance, the mitochondrial 60 kDa heat shock protein (CH60), aconitate hydratase (ACON), acyl-coenzyme A synthetase ACSM2 (ACSM2), and succinyl-CoA:3-ketoacid coenzyme A transferase 1 (SCOT1) exhibited at least 10 Sirt5-regulated lysine residues.

**Figure 5.**
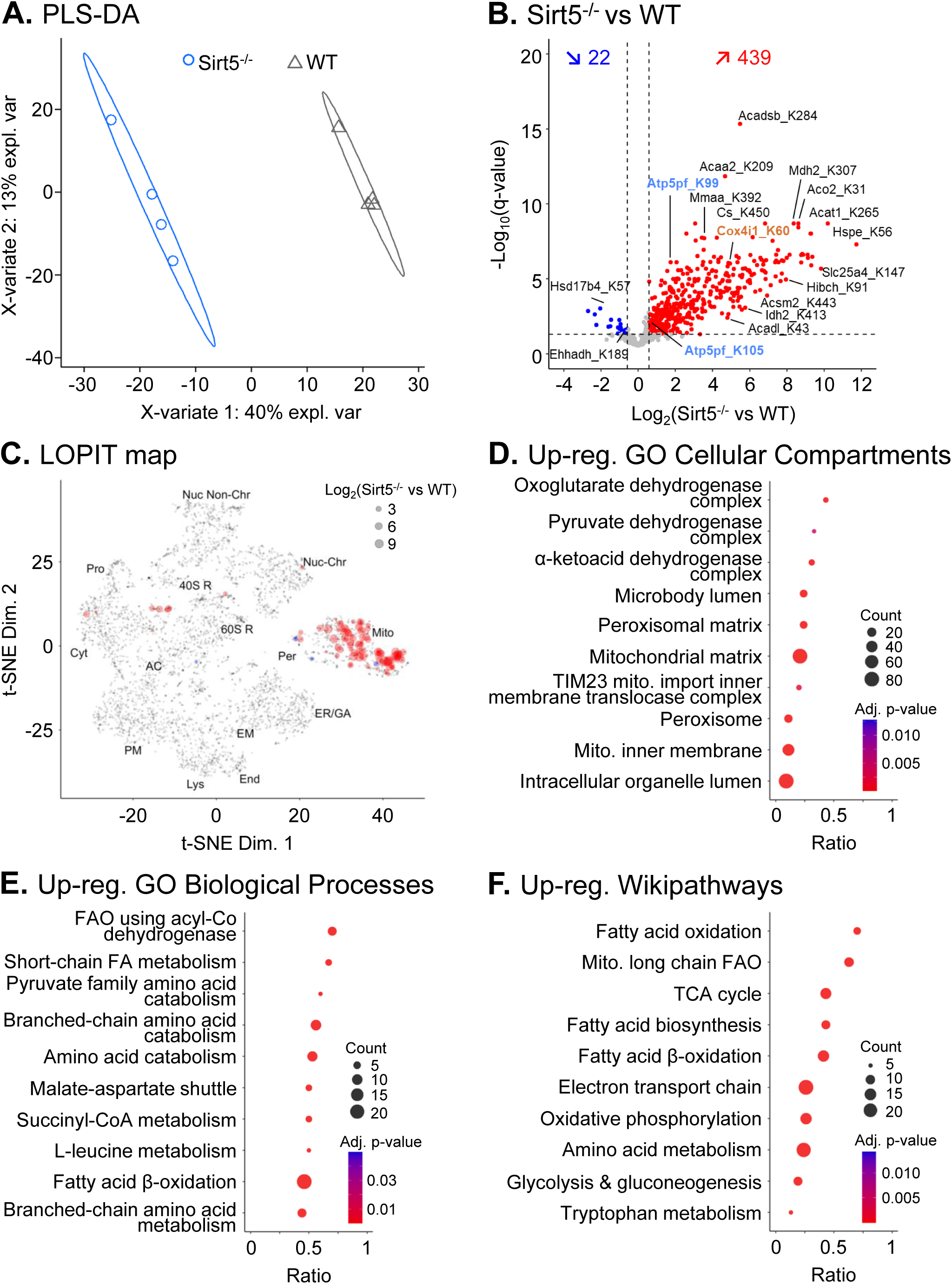
Sirt5 deletion led to a hypersuccinylation of mitochondrial proteins related to metabolism in mouse kidneys. **A.** Supervised clustering using partial least-square discriminant analysis of all succinylated peptides identified and quantified in Sirt5^-/-^ (blue dot) and WT (grey triangle) tissues. **B.** Volcano plot showing the remodeling of the 980 quantified succinylated sites in Sirt5^-/-^ vs WT mouse kidneys (q-value < 0.05 and |log_2_(fold-change)| > 0.58). **C.** LOPIT map representing the subcellular distribution of the Sirt5-targeted succinylated proteins. Dots represent the log_2_(Sirt5^-/-^ vs WT) obtained for the succinylated site with the maximum absolute fold-change for each protein, where up-regulation is displayed in red and down-regulation in blue. Note that one protein can present more than one significantly altered succinylated site. Assigned coordinated for 103 succinylated proteins for the 40S Ribosome (40S R), 60S Ribosome (60S R), Actin Cytoskeleton (AC), Cytosol (Cyt), Endoplasmic Reticulum/Golgi Apparatus (ER/GA), Endosome (End), Extracellular Matrix (EM), Lysosome (Lys), Mitochondria (Mito), Nucleus – Chromatin (Nuc-Chr), Nucleus – Non-chromatin (Nuc Non-Chr), Peroxisome (Per), Plasma Membrane (PM), Proteasome (Pro), and Undefined (grey) subcellular locations are shown, when available (based on the localization of unmodified proteins as determined in^23^). **D-F.** Dotplots representing the top10 Enrichr^25^ **D.** GO Cellular Compartments, **E.** GO Biological Processes, and **F.** Wikipathways enriched for up-regulated succinylated proteins upon Sirt5 deletion.

To gain more cellular and biological insights into the targeted succinylated proteins, we first generated an *in silico* LOPIT map against the reference mouse pluripotent stem cell subcellular proteome dataset^23^ to inform on the subcellular localization of the modified proteins (**Fig. 5C**). This revealed that the mitochondrial sub- proteome was primarily targeted by Sirt5, which aligns with the known mitochondrial location of Sirt5 in cells^24^. More specifically, a Gene Ontology (GO) cellular compartment enrichment analysis conducted in Enrichr^25^ showed that these proteins were part of the oxoglutarate dehydrogenase complex, pyruvate dehydrogenase complex, and α-ketoacid dehydrogenase complex, among others (**Fig. 5D**; **Supplementary Table 4**). Several targets were also located to the peroxisomal compartment, including the dehydrogenase/reductase SDR family member 4 (DHRS4) and the peroxisomal bifunctional enzyme (ECHP). With respect to the biological pathways, the enrichment analysis pointed to an up-regulation of GO biological processes such as fatty acid β-oxidation using acyl-CoA dehydrogenase, short-chain fatty acid metabolism, branched-chain amino acid catabolism, and succinyl-CoA metabolism (**Fig. 5E**). Up-regulated Wikipathways additionally highlighted the tricarboxylic acid (TCA) cycle, electron transport chain, and oxidative phosphorylation (**Fig. 5F**).

A more detailed view of the succinylated proteins that are targeted by Sirt5 and part of the fatty acid β-oxidation (FAO) pathway and TCA cycle are depicted on **Fig. 6**. After the transport of acyl-CoAs and fatty acids into the mitochondrial matrix and the activation of the latter by CoA to produce reactive acyl-CoA species, FAO will further lead to the shortening of the acyl-CoA (C_n_) species into acyl-CoA (C_n-2_) and the production of a molecule of acetyl-CoA via a stepwise enzymatic reaction. Multiple enzymes are involved at each of these five key steps that depend on the length of the acyl-CoA chain. Sirt5-targeted proteins identified in our study were involved in each reaction of FAO. Acetyl-CoA species, generated via FAO or enzymatic oxidation of pyruvate, can enter the TCA cycle. This eight-step process will lead to the production of reduced electron carriers (NADH and FADH_2_), that are critical to oxidative phosphorylation and thus cellular respiration. For each step of this central process as well, multiple succinylation sites of the enzyme players were targeted by Sirt5. In parallel to this, one FAO-shortened acyl-CoA product is propionyl-CoA (C_3_), that can further be converted to succinyl-CoA, ultimately entering the TCA cycle. The last step of this process, L-methylmalonyl-CoA isomerization into succinyl-CoA, is performed by methylmalonyl-CoA mutase (MMUT), which requires adenosylcobalamin, the biologically active form of vitamin B12, as a coenzyme. Interestingly, we report here that Sirt5 targets Lys-392 of the mitochondrial methylmalonic aciduria type A homolog (MMAA), a protein involved in vitamin B12 transport and processing that is essential for MMUT activity^26, 27^ (see **Supplementary** Fig. 4).

**Figure 6.**
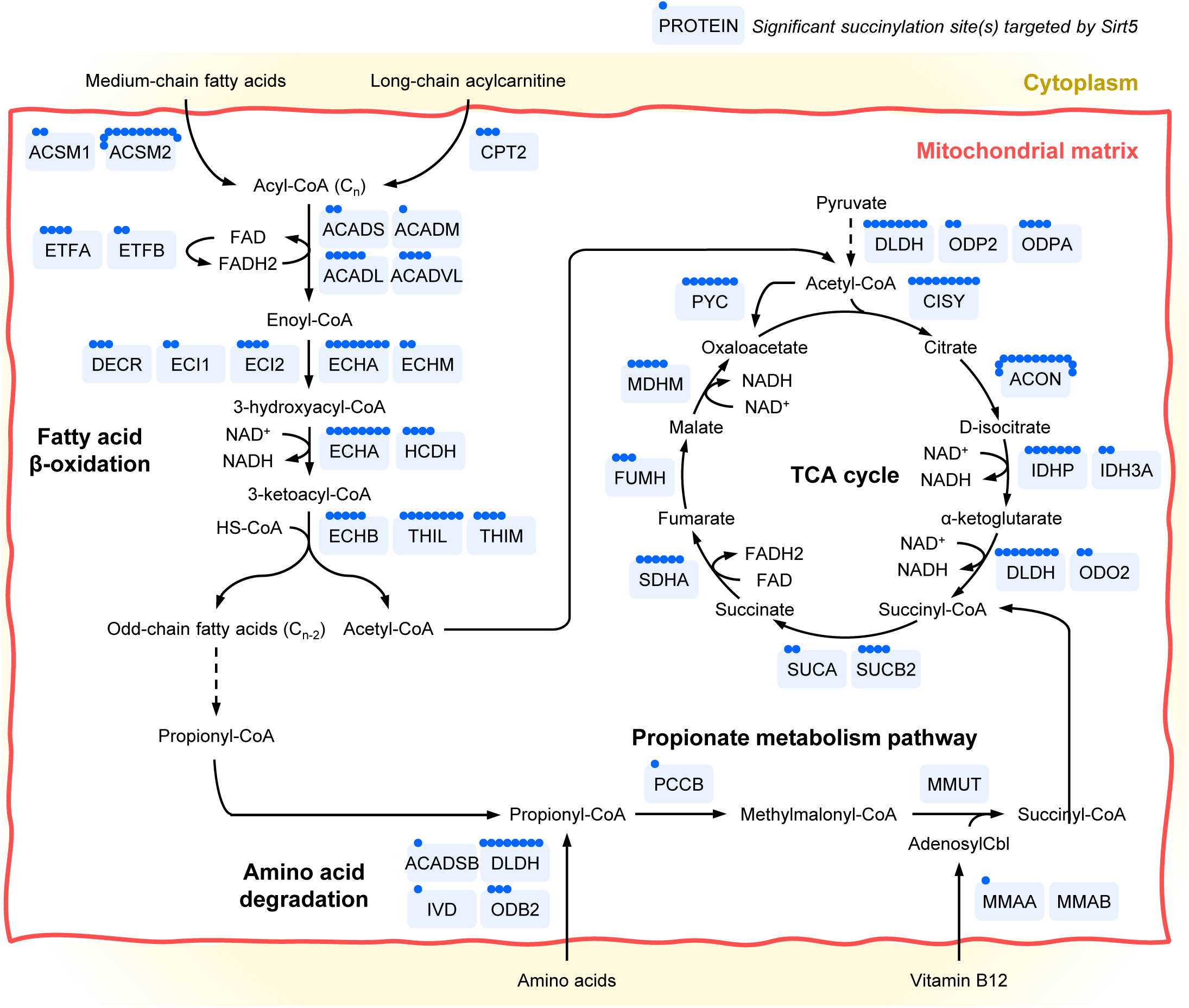
Diagram representing Sirt5-targeted mitochondrial succinylated enzymes composing the fatty acid. β**-oxidation pathway, amino acid degradation, tricarboxylic (TCA) cycle, and propionate metabolism pathway in mouse kidneys.** Each blue dot represents one significantly altered succinylation site in Sirt5^-/-^ vs WT mouse kidneys identified in this study.

## Discussion

Here, we present a novel HRIM workflow using Parallel Accumulation with Mobility-Aligned Fragmentation (PAMAF) acquisition mode for comprehensive and reproducible proteome-wide succinylome analysis. Unlike typical proteomics workflows employed for PTM analysis, based on data-dependent acquisition (DDA) or data-independent acquisition (DIA), the PAMAF strategy does not rely on the quadrupole to isolate precursor ions contained within defined mass windows. Instead, ions are, in parallel, accumulated and then separated in a 13-meter-long path traveling wave IMS module, after which low collision energy (MS1) and high collision energy (MS2) frames are alternatively acquired for all ions exiting the SLIM module. The association between the paired precursor and fragment ions is inferred based on alignment in the IM dimension, analogous to LC coelution analysis for MS1 isotopes or MS2 fragments. The current PAMAF-MS data output files can be processed by Skyline and Spectronaut. These software solutions provide easy and straightforward processing pipelines for mobility- filtered data interpretation and visualization.

The dual and orthogonal separation of proteolytic peptides using liquid chromatography and high-resolution SLIM ion mobility combined with full spectrum acquisition of all detectable precursor and fragment ions reduced complexity, thus increasing sensitivity, and provided enhanced specificity and sustained optimal ion utilization. By analyzing synthetic succinylated peptides (S1) and mouse kidney succinylomes (S2), we demonstrated that the PAMAF technology was able to fully resolve co-eluting, isobaric and isomeric succinylated peptides at the amino acid level that could not be differentiated solely in the LC and *m/z* dimensions, *i.e.* without HRIM. Importantly, it provided confident PTM site localization, as evidenced by the detection of discriminating fragment ions, even for succinylated isomer species, which is often a challenge for large-scale PTM analysis but highly relevant for downstream biological interpretations. It alleviates the need for off-line fractionation, such as high-pH reverse phase or ion exchange chromatography, to improve PTM isomer separation, which is highly advantageous when sample amounts are limited and higher throughput is preferred.

Moreover, the low cycle time of the PAMAF acquisition mode ensures that it is compatible with (ultra-)fast liquid chromatographic gradients while retaining highly quantitative performances. Indeed, enough data points across the peaks - on average, over 9 points in Study 1 (200 SPD) and over 5 points in Study 2 (15 SPD) - were collected, ensuring quantitative precision and robustness. In addition, efficient data extraction in both the LC and IM dimensions allowed for high assay specificity, reducing interferences and false identifications while improving profiling comprehensiveness, sensitivity, and quantitation.

Leveraging this novel technology, we were able to quantitatively profile hundreds of succinylation sites in kidney tissues from WT and Sirt5^-/-^ mice. As expected, Sirt5 deletion resulted in a hypersuccinylation of the mouse kidney proteome, specifically the mitochondrial sub-proteome, in line with the known subcellular localization of Sirt5 ^24^. In this organelle, Sirt5 targets are key players of a wide range of metabolic processes, spanning fatty acid β-oxidation, the TCA cycle, oxidative phosphorylation, amino acid metabolism, and ketone body metabolism, among others. While we and others have already reported similar findings^8, 10, 11, 28, 29^, this demonstrates that the novel PTM PAMAF-MS-directDIA workflow was able to capture the proteome-wide succinylome enriched from kidney tissues and accurately recapitulate its Sirt5-associated dynamic succinylome remodeling. But strikingly and to the best of our knowledge, we identified succinylated Lys-392 of the mitochondrial methylmalonic aciduria type A homolog (MMAA) as a novel target of Sirt5 in mouse kidney. In a previous and independent study on Sirt5^-/-^ mouse brain succinylome, we showed that Sirt5 regulated the succinylation of three other residues of this protein, namely Lys-78, Lys-92, and Lys-409^11^. MMAA is a GTPase involved in the metabolism of vitamin B12 by mediating the transport of cobalamin into mitochondria^27^. It also acts as a G-protein chaperone enabling the delivery of the cofactor adenosylcobalamin from MMUB to MMUT as well as exhibits dual protectase and reactivase roles for MMUT^30^. Interestingly, the lysine residue at position 392 in mouse MMAA is conserved as Lys-395 in the human ortholog, which may also be subjected to acylation events.

Beyond lysine succinylation, we believe this workflow can be applied to profile and quantify any PTMs of interest with improvements for precise quantification and site localization. We envision the use of this methodology to broad, large-scale, and high-throughput proteomics applications, ranging from, but not limited to, discovery studies to uncover molecular mechanisms underlying genetic perturbations or disease pathogenesis/progression to translational/clinical studies investigating responses to therapeutic interventions.

## Methods

### Synthetic peptides

Synthetic heavy-labeled succinylated and matched unmodified peptides are detailed in **Supplementary Table 1**. The succinylated synthetic peptides GA**^112^Ksucc**YV*SHGATGKGNDQVRFE and GA**K**YV*SHGATG**^121^Ksucc**GNDQVRFE (UniProt ID: P16460) as well as LTGA**^216^Ksucc**V*VVSGGRGLKSGE and LTGA**K**V*VVSGGRGL**^226^Ksucc**SGE (UniProt ID: Q99LC5) containing a stable isotope-labeled valine residue (V* with ^13^C_5_ and ^15^N_1_) were from New England Peptide at >99% chemical and >99% isotopic purity. Synthetic peptides were diluted in 2% acetonitrile (ACN), 0.1% formic acid (FA) in H_2_O for MS analysis.

#### Mouse kidney tissues

Age-matched 3-month-old male mice were used throughout the study. Homozygous global Sirt5 knockout mice (*Sirt5*^-/-^)^31^ were obtained from the Jackson Laboratory (B6;129-*Sirt5*^tm1Fwa^/J, Stock No. 012757). To produce appropriate littermate controls, Sirt5 heterozygous mutant mice (*Sirt5^+/–^*) were bred together to produce WT (*Sirt5^+/+^*) and (*Sirt5*^–/–^) mutants. The University of Pittsburgh Institutional Animal Care and Use Committee approved all experiments (Approval No. 22112009).

#### Sample preparation for succinylomic analysis

Kidney tissues from WT (n=4) and Sirt5^-/-^ (n=4) mice were collected. The samples were prepared for succinylomic analysis as described in previous studies^11–13^ with slight modifications. Briefly, frozen kidneys were immersed in 700 μL of lysis buffer containing 8 M urea, 200 mM triethylammonium bicarbonate (TEAB), pH 8, 75 mM sodium chloride, 1 μM trichostatin A, 3 mM nicotinamide, and 1x protease/phosphatase inhibitor cocktail (Thermo Fisher Scientific, Waltham, MA), and homogenized for 2 cycles with a Bead Beater TissueLyser II (Qiagen, Germantown, MD) at 25 Hz for 4 min each. Lysates were clarified by spinning at 15,700 x *g* for 15 min at 4°C, and the supernatant containing the soluble proteins was collected. Protein concentrations were determined using a Bicinchoninic Acid Protein (BCA) Assay (Thermo Fisher Scientific), and subsequently 3.1 mg of protein from each sample were utilized for further processing. Protein lysates were reduced using 20 mM dithiothreitol (DTT) in 50 mM TEAB for 30 min at 37 °C, and after cooling to room temperature, alkylated with 40 mM iodoacetamide (IAA) for 30 min at room temperature in the dark. Samples were diluted 10-fold with 50 mM TEAB and proteins were digested overnight with a solution of sequencing-grade trypsin (Promega, San Luis Obispo, CA) in 50 mM TEAB at a 1:50 (wt:wt) enzyme:protein ratio at 37°C. This reaction was quenched with 1% FA and the sample was clarified by centrifugation at 1,800 x *g* for 15 min at room temperature. Clarified peptide samples were desalted with Oasis 30-mg Sorbent Cartridges (Waters, Milford, MA). The proteolyzed and desalted lysates (3 mg for each sample) were re- suspended in 1.4 mL of immunoaffinity purification buffer (Cell Signaling Technology, Danvers, MA) for succinyllysine enrichment with anti-succinyl antibody conjugated to agarose beads from the Succinyl-Lysine Motif Kit (Cell Signaling Technology, Kit #13764). This process was performed according to the manufacturer protocol; however, each sample was incubated in half the recommended volume of washed beads. Peptides were eluted from the antibody-bead conjugates with 0.15% trifluoroacetic acid (TFA) in H_2_O and were desalted using C_18_ stagetips made in-ho9use. Samples were vacuum dried and re-suspended in 20 μL of 0.2% FA in water plus 1 μL of indexed retention time standard peptides (iRT; Biognosys, Schlieren, Switzerland)^32^ prepared according to manufacturer’s instructions.

#### HPLC-PAMAF-MS analysis

HPLC-PAMAF-MS acquisitions were performed on an Evosep One system coupled to an Agilent 6545 Quadrupole Time-of-Flight (Q-TOF) mass spectrometer. Synthetic peptides (500 pg on-column for each peptide) were separated using the Evosep 200 samples per day (SPD) method on an EV1107 column (4 cm x 150 µm, 1.9 µm C_18_ beads). Mouse kidney succinyl enrichments (10 μL) were acquired using the Evosep Extended 15 SPD method on the EV1137 column heated to 40°C (15 cm x 150 µm, 1.5 µm C_18_ beads). After ultra-high pressure liquid chromatography (UHPLC) separation, samples were ionized using a custom nano electrospray ionization source, with +1.85 kV at the entrance capillary. Ions were transferred through a heated glass capillary (18-cm long, 0.6 mm ID, 280°C) into the HRIM module for ion mobility separation using a 13-m-long ion path (long path) in the SLIM module, after which they were released into the Agilent 6545 Q-TOF mass spectrometer for mass spectrometric analysis. For all mass analyses, the scan range covered 50-1700 Th. For MS2 scans, all precursor ions were transferred to the Q-TOF collision cell without any quadrupole filtering, and all ions were fragmented using a collision energy varied from 12 to 42 V depending on *m/z*. Accumulation times were set at 100 ms for MS1 and MS2 mobility frames, and cycle time was 810 ms.

#### Data analysis with Skyline – Synthetic succinylated peptides

Synthetic peptide data (*.mbi format) was processed in Skyline^21^ (Skyline-daily, version 25.0.9.131). A study-specific ion mobility library was generated from a representative acquisition file and library arrival time values were manually adjusted, if needed (**Supplementary Table 2**; for step-by-step protocol, please see Dodds *et al.*^22^). The IM library was used for mobility filtering of the data, applying a resolving power set at 50. For MS1 and MS/MS filtering, resolving power was set to 20,000. DIA was selected as acquisition method and all ions as isolation scheme. All matching scans were used. Product ions (mono- and doubly-charged y- and b-type ions) from ion-2 to last ion-1 were extracted. Chromatographic peaks were manually checked to remove interfered transitions and correct peak boundaries.

#### Data analysis with directDIA (Spectronaut) – Succinylation enrichments from mouse kidneys

Mouse kidney succinylated peptide enrichment acquisition files in *.mbi format were first converted to a *.bgms (Biognosysis generic MS) format using a currently proprietary tool. Data were processed in Spectronaut (version 19.4.241104.62635) using directDIA. Data were searched against the *Mus musculus* reference proteome with 54,910 protein entries (UniProtKB-TrEMBL), accessed on 05/18/2024. Trypsin/P was set as digestion enzyme, and two missed cleavages were allowed. Cysteine carbamidomethylation was set as fixed modification, and lysine succinylation, methionine oxidation and protein N-terminus acetylation as variable modifications. Data extraction parameters were set as dynamic. Identification was performed using 1% precursor and protein q-value. The PTM localization score was selected with a probability cutoff of 0.75. Quantification was based on the extracted ion chromatograms (XICs) of 3–6 MS2 fragment ions, specifically b- and y-ions, without normalization, as well as data filtering with q-value sparse. Grouping and quantitation of PTM peptides were accomplished using the following criteria: minor grouping by modified sequence and minor group quantity by mean precursor quantity. Differential expression analysis was performed using an unpaired t-test, and p-values were corrected for multiple testing using the Storey method^33, 34^. For determining differential changes, a q-value < 0.05 and absolute log_2_(fold-change) > 0.58 was required (**Supplementary Table 3**).

#### Clustering analysis

Partial least squares-discriminant analysis was performed on the mouse kidney succinylated peptide data using the mixOmics package^35^ in RStudio (version 2025.05.1; R version 4.5.1).

#### LOPIT organellar protein distribution

The hyperLOPIT2015 pluripotent stem cell dataset was downloaded in Rstudio using the pRolocData R package (Bioconductor) and plotted with the pRoloc R package^36^. To create the reference LOPIT organellar protein distribution dataset, a crude membrane preparation from mouse pluripotent stem cell lysate was fractionated by density gradient ultracentrifugation to separate and enrich organelles based on density^23^. Proteins were identified via quantitative multiplexed MS and assigned to organelles based on similarities in distribution in the density gradient to well-annotated organelle protein markers^37, 38^. The t-SNE machine learning algorithm was used to reduce the number of dimensions in the LOPIT proteomic data to a 2D map where proteins cluster by similarity from multiple experimental factors^39, 40^. All LOPIT protein assignments were used as originally determined and without additional refinement.

#### Enrichment analysis

Up-regulated succinylated genes were imputed into Enrichr^25^, a web-based tool for enrichment analysis, using all identified and quantified genes as background. Identified Gene Ontology (GO) terms and pathways were filtered for an adjusted p-value < 0.05 (**Supplementary Table 4**). Dotplots were generated using the ggplot2 package^41^ in RStudio.

## Data Availability

The mass spectrometry data that support the findings of this study are available via the FTP link: ftp://MSV000099489@massive-ftp.ucsd.edu or via the MassIVE website: https://massive.ucsd.edu/ProteoSAFe/dataset.jsp?task=5148936c9fc0440a9adba9f20d c70fee (MassIVE ID: MSV000099489; ProteomeXchange ID: PXD069456).

[Note to the reviewers: To access the data repository MassIVE (UCSD) for MS data, please use: Username: MSV000099489_reviewer; Password: winter].

## Acknowledgments

We thank Eleni Vickers for help with the initial data processing. This work was supported by NIDDK R01 DK121758 and R01 DK134346 and by R25 DK119180 (S.S.L.).

## Author Contributions

B.S. contributed to study design, conceptual development, data interpretation, manuscript revision and editing, and provided resources. L.R.III contributed to data acquisition and analysis, manuscript and figure preparation, and manuscript editing.

L.R. contributed to data analysis and manuscript editing. A.S.B. and K.P. contributed to murine sample collection, and manuscript editing. E.G. and S.S.L. contributed to study design and manuscript editing. D.D. contributed to study design, conceptual development, manuscript editing, and provided resources. J.B. contributed to study design, conceptual development, sample preparation, data interpretation, manuscript and figure preparation and revision.

## Competing Interests

B.S. serves on the proteomics advisory board for MOBILion Systems, Inc.. L.R. III, L.R., and D.D. are employees of MOBILion Systems, Inc.. The other authors declare no competing financial interest.

---

**Supplementary Figure 1.** PAMAF mode combined with ion mobility-filtering in Skyline for confident succinylated isomer differentiation. The synthetic succinylated peptides LTGA^216^KsuccV*VVSGGRGLKSGE (*top*) and LTGAKV*VVSGGRGL^226^KsuccSGE (*bottom*) from the mouse mitochondrial electron transfer flavoprotein subunit alpha (UniProt ID: Q99LC5) were acquired on the prototype SLIM-based HRIM Q-TOF (Agilent 6545) MS operated in PAMAF mode. Data were analyzed in Skyline using an ion mobility library. Ion mobility-filtered A. MS/MS spectra and B. extracted ion chromatograms (XIC) are displayed for the two succinylated peptides (precursor ions at *m/z* 607.67, z = 3) and show the detection of strong differentiating fragment ions for precise PTM localization. This demonstrates that the PAMAF mode can differentiate positional isomers with high confidence and sensitivity performance compatible with quantification.

**Supplementary Figure 2.** MS/MS spectra of HRIM-PAMAF-resolved co-fragmented species. The overlapping isotope distributions in succinylated peptide enrichments from Sirt5^-/-^ and WT mouse kidneys displayed on Figure 4 were resolved by HRIM achieved with the PAMAF technology. The corresponding ion mobility-filtered MS/MS spectra (RT 64.44 – 64.82 min) are displayed and the identification information obtained from the directDIA search are mentioned. For the succinylated peptide DYPLPDVAHVTMoxLSASQ**^60^Ksucc**ALK of the mitochondrial cytochrome c oxidase subunit 4 isoform 1 (Cox4i1) with a precursor ion at m/z 800.74 and AT 246.11 ms, several fragment ions containing the succinyl group were detected, enabling precise and confident PTM site localization.

**Supplementary Figure 3.** Representative extracted ion chromatogram of the precursor ions at m/z 882.92 in a succinylated peptide enrichment of Sirt5^-/-^ mouse kidney. Two species were detected at the MS1 level and further identified using ion mobility-resolved MS/MS using PAMAF technology: FDDPKFEVID^105^KsuccPQS and FDDP^99^KsuccFEVIDKPQS from the mouse mitochondrial ATP synthase-coupling factor 6 (Atp5pf) that eluted at 56.9 min and 57.5 min, respectively.

**Supplementary Figure 4.** The Lys-392 of the mitochondrial methylmalonic aciduria type A homolog is succinylated in mouse kidney. **A.** Representative MS1 spectrum, mobilogram and heatmap of the precursor ion at *m/z* 592.835 (z = 2) and arrival time 196.72 ms obtained in a succinylated peptide enrichment of Sirt5^-/-^ mouse kidney analyzed in PAMAF mode. **B.** The corresponding MS/MS spectrum led to the identification of the succinylated peptide sequence **^392^Ksucc**VLSGALSPGR from the mouse mitochondrial methylmalonic aciduria type A homolog (Mmaa). The detection of succinyl-containing fragment ions, especially the ions b_2_ and b_3_ (in red), provides strong evidence for the confident localization of the succinyl group on Lys-392. **C.** The 3D structure of the mitochondrial methylmalonic aciduria type A homolog (UniProt ID: Q8C7H1; predicted structure AlphaFold AF-Q8C7H1-F1) was viewed with the PTMVision web server for PTM visualization from proteomics data^42^. The zoom in the protein 3D structure (*right*) shows the distance of the succinylated residue Lys-392 to the neighboring residue Ala-397 (4.54 Å) and Leu-398 (3.92 Å). This proximity could lead to the interaction between the negatively charged succinylation group and the neighboring alanine and leucine residues and thus change the proteoform conformation. This hypothesis together with the succinylation site validation and its functional properties need to be verified and assessed in follow-up studies.

