## Supplementary figures and images for "Novel high-resolution ion mobility mass spectrometry for site-specific quantification of the sirtuin-5 regulated kidney succinylome"

### Supplemental Figure 1

#### Supplementary Figure 1

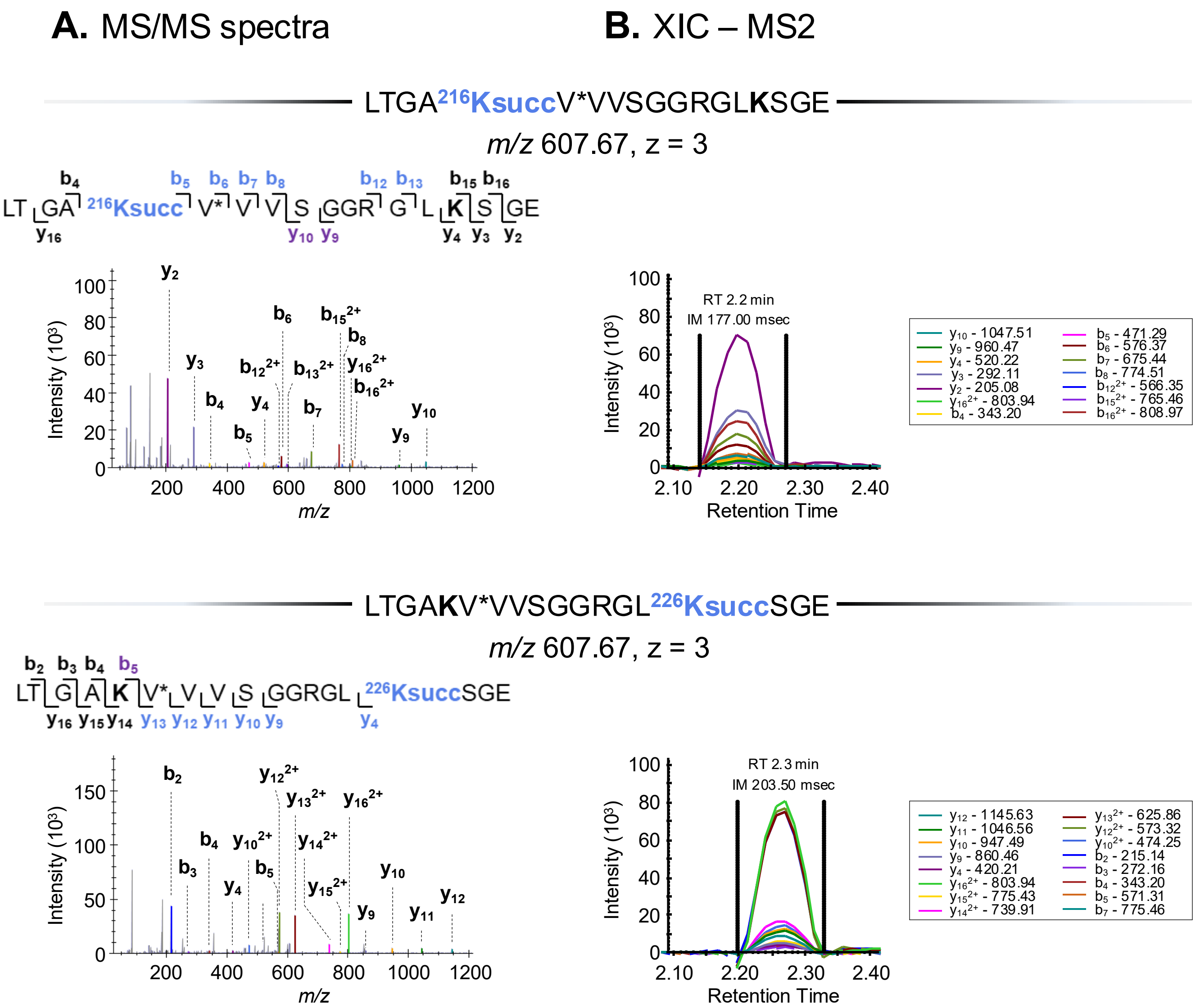

### Supplemental Figure 2

#### Supplementary Figure 2

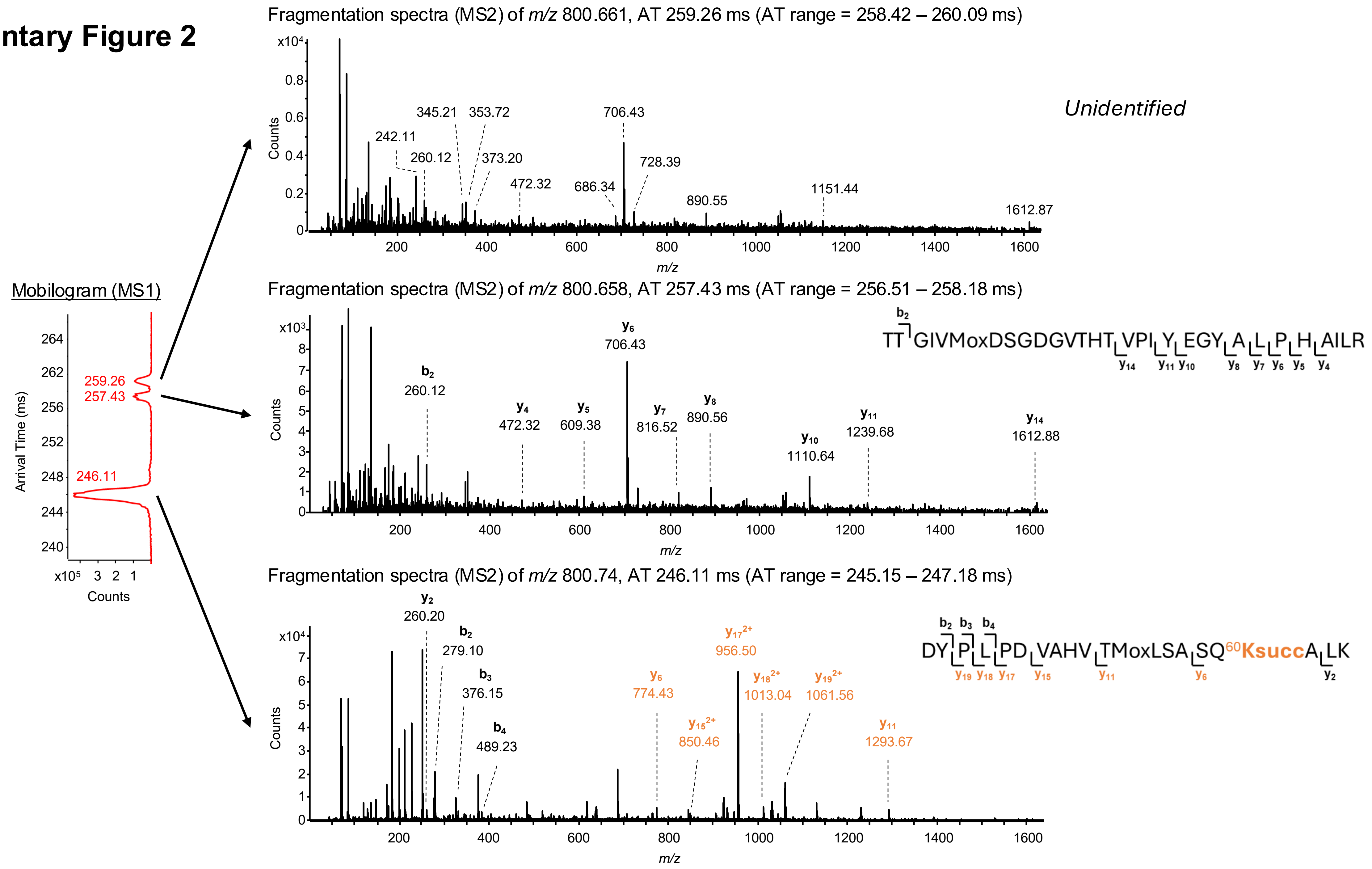

### Supplemental Figure 3

#### Supplementary Figure 3

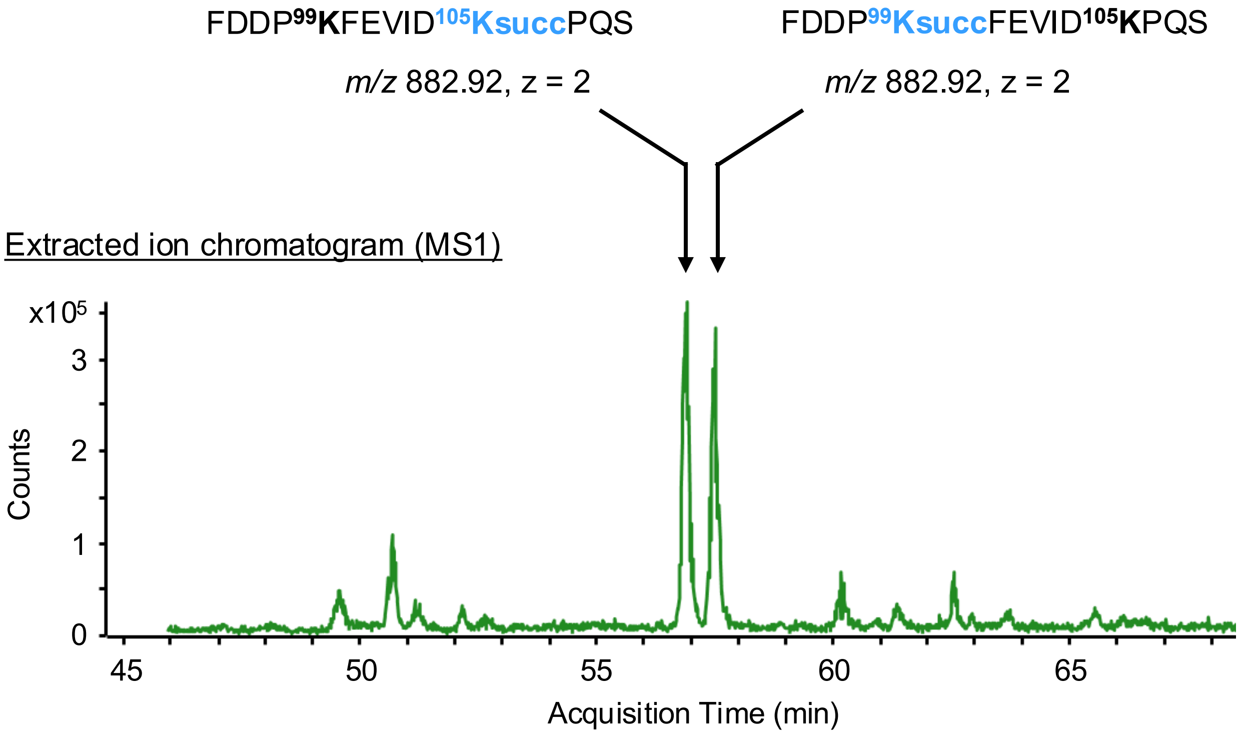
