## Supplemental Figure 4 for "Novel high-resolution ion mobility mass spectrometry for site-specific quantification of the sirtuin-5 regulated kidney succinylome"

### Supplementary Figure 4

#### A. MS1 spectrum and mobilogram

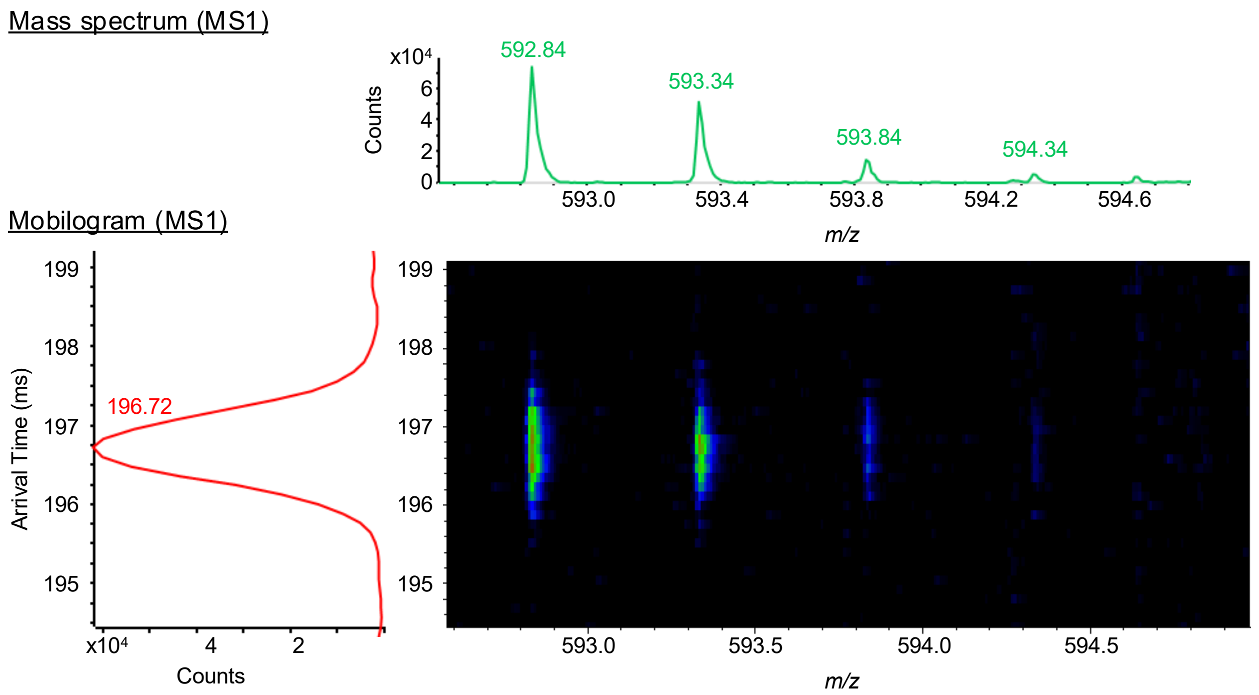


#### B. MS/MS spectrum – *m/z* 592.835; z = 2

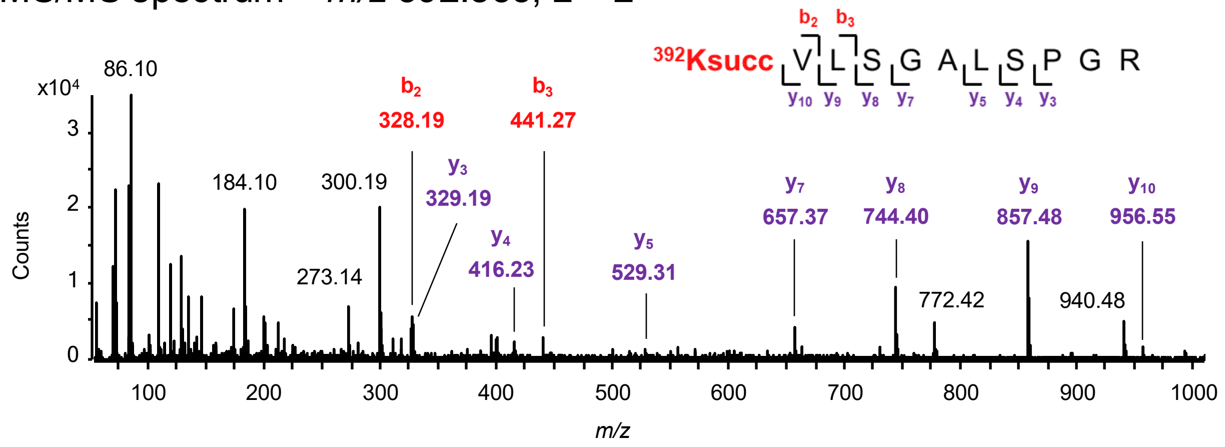


#### C. Structure and succinylated Lys-392 potential residue contacts

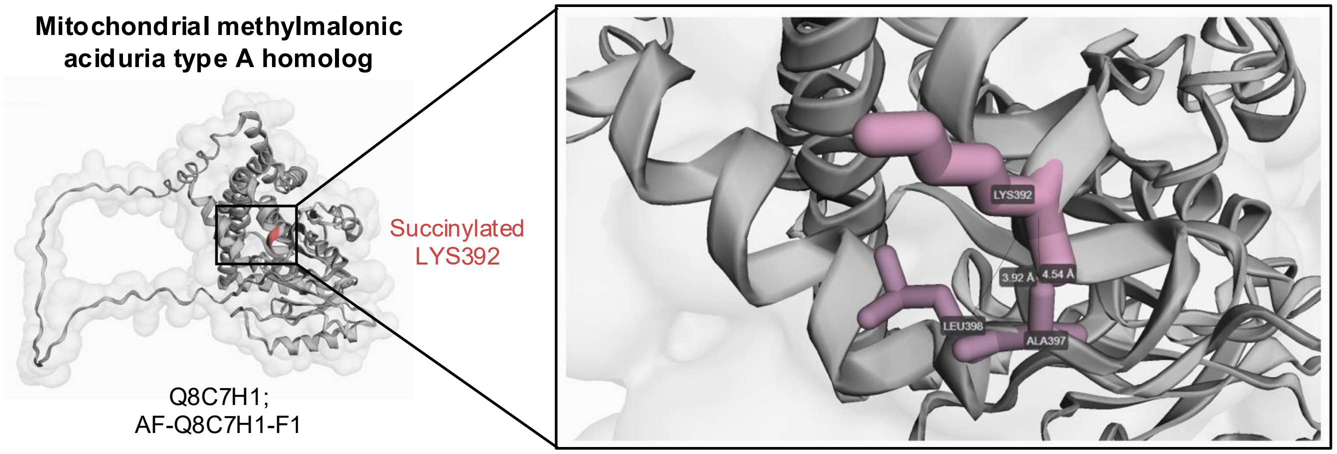
